# Clonal effects of the *Ras* oncogene revealed by somatic mutagenesis in a *Drosophila* cancer model

**DOI:** 10.1101/2025.05.08.652841

**Authors:** Takuya Akiyama, Matthew C. Gibson

**Author notes:** Corresponding authors Takuya Akiyama, Matthew C. Gibson.

## Abstract

Somatic mutations of *Ras*, encoding a small GTPase, are detected in a wide range of human cancers. Tumor genome sequencing further reveals a cancer type-dependent mutational spectrum for the *Ras* gene, suggesting that tissue- and allele-specific effects underlie tumorigenic activity. Although biochemical studies have characterized the GTPase activity of several Ras variants *in vitro*, precisely how somatic mutations of the endogenous *Ras* locus differentially affect tissue growth and homeostasis remain elusive. Here we engineered the endogenous *Drosophila Ras* locus to create a spectrum of inducible oncogenic alleles and then compared their activities *in vivo*. In the developing wing primordium, somatic clones carrying the oncogenic mutation *Ras G12V* exhibited a weak activation of downstream MAPK signaling but did not disrupt tissue architecture. However, cell clones carrying the same *Ras G12V* allele in the adult midgut exhibited a growth advantage and progressively took over the tissue, resulting in intestinal barrier dysfunction. In contrast, cell clones expressing a distinct allele, *Ras Q61H*, formed aberrant cysts that disrupted epithelial architecture and triggered local cell death. Conversely, when we induced cell clones carrying *Ras Q61H* in the midgut, hyper-proliferating mutant cells rapidly expanded to occupy the entire tissue. Surprisingly, this population of rapidly expanding mutant cells was eventually eliminated from the midgut, restoring *wild-type* cells and normal barrier function. Thus, in the midgut, *Ras G12V* was ultimately more deleterious than *Ras Q61H* due to the regression of *Ras Q61H* mutant cells. These results establish a new model for somatic mutagenesis at the *Ras* locus and illuminate a mechanistic basis for the tissue-specific effects of oncogenic *Ras* variants. Further, this study provides direct evidence that allele-dependent clonal dynamics may play a critical role in the tissue-selectivity of *Ras* oncogenic mutations.

## Introduction

Carcinomas, which originate from epithelial tissues, are the most common type of cancer and are responsible for over 80% of cancer-related deaths in the Western world^1^. Oncogenic mutations in the *Ras* gene are pervasive throughout various types of cancer, including colorectal and lung cancers, and are responsible for approximately 20% of all cancer cases^2, 3^. *Ras* was discovered in the 1960s through studies of a retrovirus that induces rat sarcoma^4, 5^. In the 1980s, *Ras* was identified as the first human oncogene isolated from a bladder carcinoma cell line^6–8^. The *Ras* gene encodes a highly conserved small GTPase consisting of several protein domains, including the P-loop and switch-I and II regions (Figure 1A; Figure S1). The human genome contains three *Ras* family genes: *K-Ras*, *H-Ras*, and *N-Ras*, while *Drosophila* has a single *Ras* gene, *Ras85D* (Figure S1).

**Figure 1.**
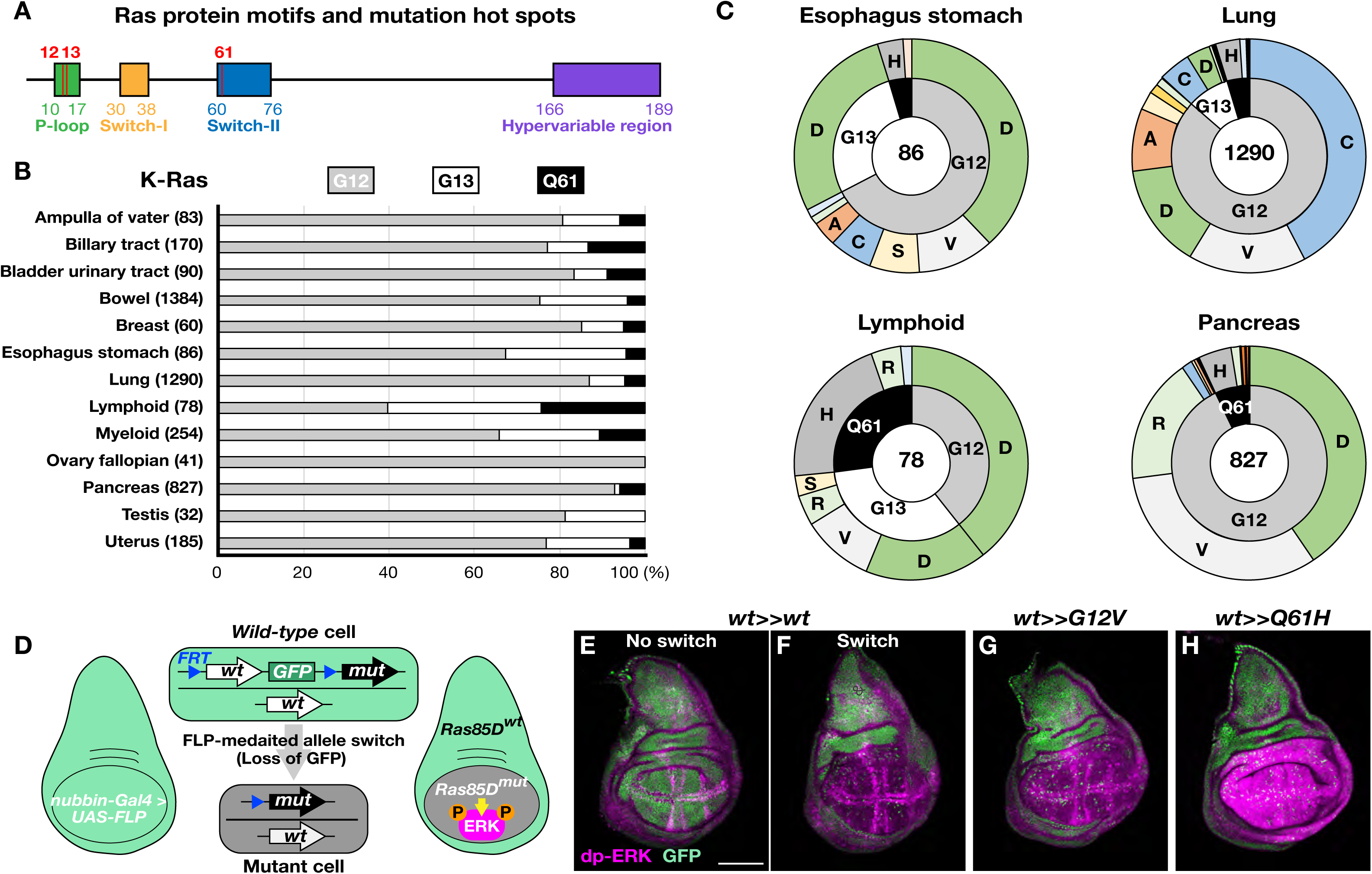
Inducible *RAS* oncogenic alleles. (**A**) Ras proteins consist of 189 amino acids and are highly conserved between organisms. They contain four protein motifs: p-loop, switch-I, switch-II, and hypervariable regions. *Ras* oncogenic mutations are mainly found at glycine 12 (G12), glycine 13 (G13), and glutamine 61 (Q61). (**B**) Tissue-specific accumulation of *K-Ras* oncogenic mutations at G12, G13 and Q61. The numbers in parentheses indicate sample sizes of *K-Ras* mutations. (**C**) Tissue-selectivity of *K-Ras* oncogenic mutation alleles. The sample sizes are shown in the center of the charts. (**D**) Induction of endogenous *Ras* oncogenic mutations via FLP/*FRT*-mediated recombination using a wing pouch-specific Gal4 driver, *nubbin-Ga4*. (**E**-**H**) Control wing discs before (**E**, *w; Ras85D^GFP>>wt^/+*) and after the *Ras85D* allele switch (**F**, *w; nubbin-Gal4/+; Ras85D^GFP>>wt^/UAS-FLP*). Induction of *Ras G12V* (**G**, *w; nubbin-Gal4/+; Ras85D^GFP>>G12V^/UAS-FLP*) and *Q61H* oncogenic mutations (**H**, *w; nubbin-Gal4/+; Ras85D^GFP>>Q61H^/UAS-FLP*) in the wing discs. Loss of GFP (g*reen*) signals indicate cells expressing desired *Ras85D* alleles due to the allele switch. dp-ERK antibody is used to monitor Ras/MAPK pathway activity (*magenta*). Scale bar,100 μm.

Ras GTPases play crucial roles in regulating cell growth, proliferation, differentiation, and cytoskeleton remodeling by controlling several downstream signaling pathways, such as the mitogen-activated protein kinase (MAPK) signaling pathway^9–11^. As aberrant activity is associated with cancer, Ras activity must be tightly controlled by alternating between its active GTP-bound and inactive GDP-bound forms^9–11^. Conversion from the active to the inactive states occurs through GTP hydrolysis, a process stimulated by GTPase-activating proteins (GAPs)^12^. When Ras is in its active state, the switch-I and switch-II regions undergo conformational changes that enable the Ras protein to interact with effector proteins to activate the downstream signaling cascades^13^. Most oncogenic mutations in *K-Ras*, *H-Ras*, and *N-Ras* are missense mutations occurring at glycine position 12 (G12), glycine position 13 (G13), and glutamine position 61 (Q61) (Figure 1A; Figure S1)^2, 3^. These oncogenic mutations prevent GAP function, leaving Ras in its active GTP-bound state^14^.

Theoretically, if the development of *Ras*-driven cancer relied solely on its oncogenic activity, *Ras* mutations with stronger oncogenic activity would be more common across all cancer types compared to those with weaker activity. Nevertheless, cancer genome sequencing has revealed that distinct *Ras* oncogenic mutations accumulate in a tissue-specific manner (Figure 1B and C; Figure S2; cBioPortal for Cancer Genomics (https://www.cbioportal.org/))^2, 3^. Given that somatic mutations occur spontaneously during development and homeostasis, and that the likelihood of acquiring particular *Ras* oncogenic mutations should be similar in each tissue^15–19^, these results suggest that selective effects within tissues play a critical role in the selectivity of different *Ras* mutations. Understanding these mechanisms could offer new insights into how *Ras* oncogenic mutations contribute to cancer initiation and development.

## Results

### *Ras* oncogenic mutations activate MAPK signaling in an allele-dependent manner

To systemically investigate the effects of distinct *Ras* oncogenic mutations under physiological conditions in *Drosophila*, we used CRISPR-Cas9 genome editing system to modify the endogenous *Ras85D* locus and establish several inducible mutant lines: *Ras85D^GFP>>wt^*, *Ras85D^GFP>>G12D^, Ras85D^GFP>>G12V^*, *Ras85D^GFP>>G13D^*, *Ras85D^GFP>>G13R^*, and *Ras85D^GFP>>Q61H^* (*Ras85D* allele switch lines; Figure S3A). We selected the larval wing imaginal disc as an initial system for phenotypic analysis, as previous studies have examined *Ras* activity in this context^20–22^. Indeed, extensive prior work has established a conceptual framework for examining the activity of oncogenes and tumor suppressors within the relatively simple epithelial context of the wing disc^23, 24^. To permit inducibility of oncogenic mutations within the endogenous *Ras85D* locus, each allele switch line contained two *flippase recognition targets* (*FRT*s), one located between exons 1 and 2 and one after the GFP marker. By default, these modified loci expressed *wild-type Ras85D* together with ubiquitous GFP expression (Figure 1D and E; Figure S3B, E, H, K, N, and Q). Upon induction, however, Flippase (FLP)-mediated recombination removes the *Ras85D* genomic region between the two *FRTs*, resulting in the loss of GFP and the insertion of modified *Ras* alleles under the control of the endogenous gene regulatory architecture (Figure 1D-H; Figure S3B-S).

We first used the Gal4/UAS system to trigger FLP-mediated recombination specifically in the wing pouch area to induce the expression of *Ras* oncogenic alleles^25^. To verify Ras activity *in vivo*, we visualized MAPK activity with an anti-phospho-p44/42 MAPK (dp-ERK) antibody. As expected, in the absence of FLP induction wing discs from all inducible lines displayed the known pattern of MAPK activity in the future longitudinal L3, L4, and L5 veins, as well as in the marginal veins along the dorsoventral boundary, consistent with previous reports (Figure 1E; Figure S3B, E, H, K, N, and Q)^26^.

We next performed control experiments to test the effect of replacing endogenous *Ras85D* with a *wild-type* copy of itself. As predicted, *nubbin-Gal4/+; Ras85D^GFP>>wt^/UAS-FLP* wing discs demonstrated highly efficient FLP-mediated recombination while maintaining MAPK activity patterns indistinguishable from *wild-type* controls (Figure 1F; Figure S3C and D). In contrast, each FLP-induced *Ras85D* oncogenic allele induced downstream MAPK activity at distinct levels (Figure 1F-H; Figure S3B-S). Among the oncogenic alleles we induced in the wing disc, *Ras G12V* mutant cells exhibited the weakest effect on MAPK activity while *Ras Q61H* elicited the most potent activation (Figure 1G and H; Figure S3H, I, J, Q, R, and S). In contrast with Gal4/UAS-mediated gene misexpression approaches, these experiments establish the *Ras85D* allele switch system as a new paradigm for the precise induction of mutations within endogenous loci, allowing for the analysis of oncogenic activity under physiological conditions.

### *Ras Q61H* triggers clone-autonomous cystogenesis in the developing wing disc

*Ras* oncogenic mutations are thought to contribute to tumor development by activating downstream signaling pathways that promote cell proliferation and survival^27, 28^. We therefore tested the effects of allele switch-induced *Ras G12V* and *Q61H* expression on cell proliferation in developing wing discs (Figure 2A-D). After inducing each oncogenic allele in the wing pouch area (using *nubbin-Gal4*>*UAS-FLP*), mitotic cells were labelled with anti-phospho-Histone H3 (p-H3). Consistent with previous findings, we found that neither *Ras G12V* nor *Q61H-*expressing cells exhibited obvious effects on cell proliferation (Figure 2A-D)^22^.

**Figure 2.**
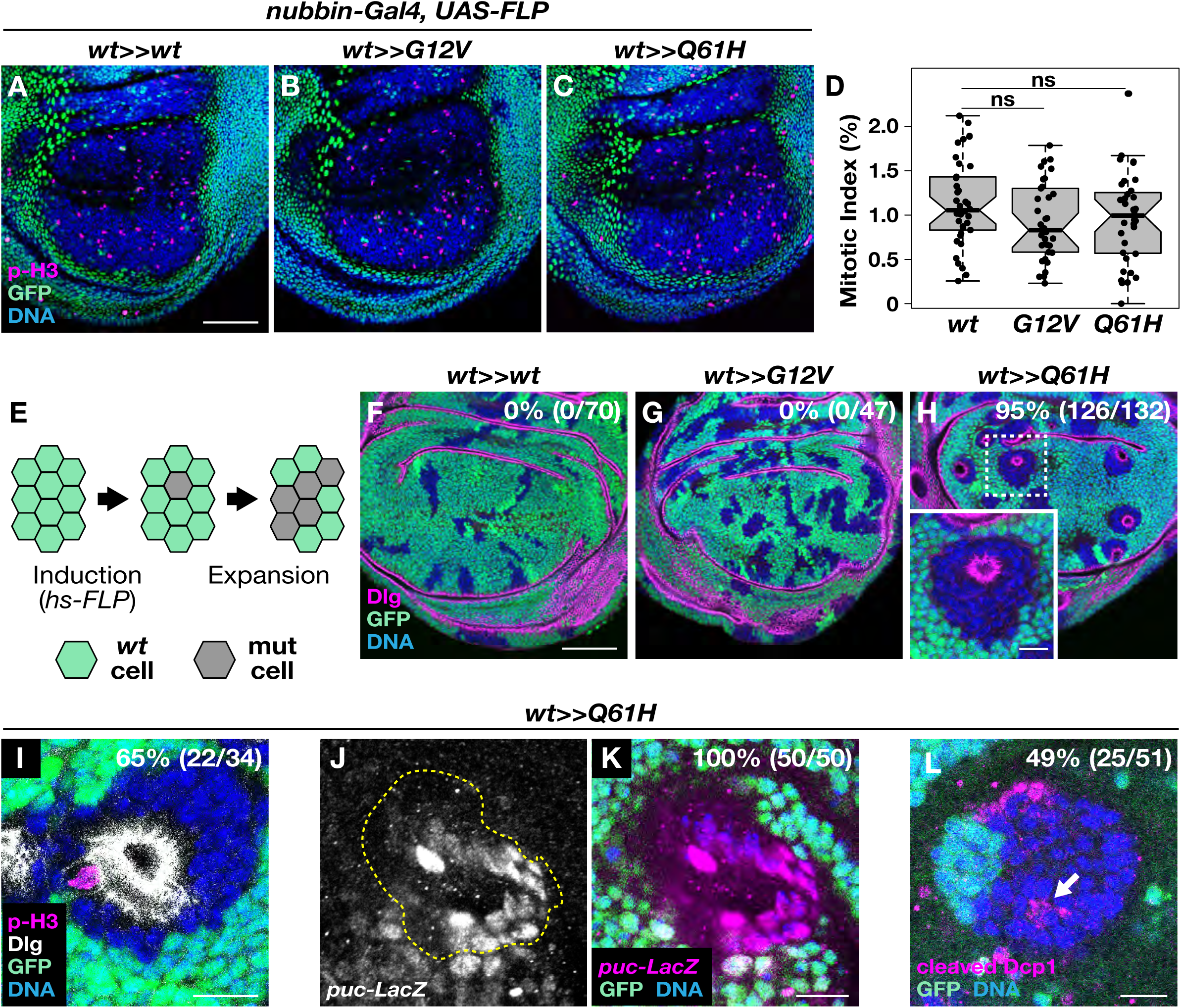
*Ras Q61H* mutation induces cysts in the developing wing disc. (**A**-**C**) *Ras* oncogenic mutations do not increase cell proliferation. Both inducing *Ras G12V* and *Q61H* mutations in the wing pouch area using *nubbin-Gal4* and *UAS-FLP* (no GFP cells in **B**, **C**) exhibit a similar number of dividing cells compared to the control wing disc (**A**). Mitotic cells are visualized by p-H3 antibody. (**D**) Mitotic index of *nubbin-Gal4/+; Ras85D^GFP>>wt^/UAS-FLP* control (*n*=38), *nubbin-Gal4/+; Ras85D^GFP>>G12V^/UAS-FLP* (*n*=34) and *nubbin-Gal4/+; Ras85D^GFP>>Q61H^/UAS-FLP* (*n*=35). The two-sided Student’s t-test shows no significant difference between samples (ns; *P* > 0.05). (**E**) Random induction of *Ras* oncogenic mutations by heat shock using *hs-FLP*. (**F**-**H**) Wing discs carrying *Ras G12V* mutant cells (**G**) have a similar epithelial morphology as control discs (**F**). In contrast, inducing *Ras Q61H* mutation causes cysts, although they maintain normal apicobasal polarity (**H**). Dlg staining indicates the apical side of the epithelium. (**I**-**L**) Although dividing *Ras Q61H* mutant cells are often observed in cysts (**I**), stress response and cell death are detected in cysts and neighboring *wild-type* cells by using a *puc-lacZ* reporter and anti-cleaved-DcpI antibody staining, respectively (**J**-**L**). Scale bars, 50 μm in **A** and **F**; 10 μm in **H**, **I**, **K**, and **L**.

In a disease context, deleterious mutations arise stochastically in individual cells and not broadly throughout a tissue. To model the spontaneous generation of cells carrying a single *Ras* oncogenic allele from somatic mutagenesis, we randomly and sporadically induced *Ras* allele switching in the wing disc during the second instar larval stage using heat shock-controlled FLP (*hs-FLP*; Figure 2E). Two days after generating mutant cells, wing discs were dissected to examine the effects of each *Ras* oncogenic mutation. In control *hs-FLP/+; Ras85D^GFP>>wt^/+* wing discs, we confirmed that a 10-minute heat shock efficiently triggered sporadic allele switching, marked by the loss of GFP (Figure 2F). Cell clones expressing the *wild-type Ras85D* allele exhibited a typical elongated wedge shape along the proximal-distal axis of the wing pouch and did not disrupt the architecture of the epithelial tissue (*n*=0/70; Figure 2F). Under identical conditions, wing discs from *hs-FLP/+; Ras85D^GFP>>G12V^/+* larvae maintained normal epithelial architecture (*n*=0/47), although mutant clones were mildly rounded (Figure 2G). In contrast, cell clones expressing *Ras Q61H* adopted a dramatic cyst-like morphology, minimized their contact with neighboring *wild-type* cells, and became partially extruded from the disc epithelium (Figure 2H; *n*=126/132). As similar effects were not observed following ubiquitous allele switching under the control of *nubbin-Gal4* (Figure 2A-C), these findings suggest that clonal cell dynamics play a key role in *Ras* phenotypes and demonstrate the crucial importance of modeling oncogenic mutant effects in a mosaic context.

In developing wing discs, mitotic nuclei migrate towards the apical side of the epithelium where they undergo cell rounding and chromosome segregation, a process known as interkinetic nuclear migration^29^. We frequently observed mitotic figures at the apical pole of cells within aberrant *Ras Q61H*-expressing cysts, suggesting that the mutant cells maintain proper apicobasal polarity (Figure 2I). Supporting this idea, we found that the polarity protein Discs large (Dlg) localized correctly to the apical region of the mutant epithelia (Figure 2H). However, we also observed that the stress-signaling Jun-N-terminal kinase (JNK) pathway was activated in *Ras Q61H*-expressing clones and detected increased apoptosis at the boundary between mutant cells and their *wild-type* neighbors (Figure 2J-L). In summary, while *Ras G12V* cell clones maintained normal epithelial architecture, those expressing *Ras Q61H* exhibited tissue extrusion and aberrant cyst formation associated with cell death. Unexpectedly, after randomly inducing the *Ras85D* allele switch, we occasionally observed elevated GFP signal adjacent to GFP-negative cells in all genotypes. This could be a result of feedback regulation at the *Ras* locus. However, we did not find any biological consequences, as we confirmed the strictly cell-autonomous activation of MAPK signaling in *Ras* oncogenic mutants (Figure S4). Importantly, in addition to the results above, we also found that the *Ras* oncogenic alleles *G12D*, *G13D*, and *G13R* were all similarly capable of inducing cysts (Figure S5).

### Induction of *Ras G12V* mutant cells disrupts the intestinal epithelial barrier

We next utilized the *Drosophila* adult midgut to investigate the effects of *Ras* oncogenic mutations in a distinct tissue context. The *Drosophila* midgut is analogous to the small intestine in mammals and contains similar cell types, including intestinal stem cells (ISCs)^30–34^. These ISCs can differentiate into two functional intestinal cells, secretory enteroendocrine cells (EEs) and absorptive enterocytes (ECs), via intermediate enteroendocrine progenitors (EEPs) and enteroblasts (EBs), respectively (Figure 3A). To introduce *G12V* and *Q61H Ras* oncogenic mutations in cells of the adult midgut, we employed the *esg^ts^FLP* recombination system, which consists of *esg-Gal4, UAS-FLP, and tubulin-Gal80^ts^* (Figure 3B)^35^. Briefly, experimental animals were reared at 18°C during development to suppress *esg-Gal4* activity through the action of Gal80^ts^. We then transferred 2-day-old adult flies to 29°C for 2 days. This temperature shift temporarily lifted Gal80-mediated repression, allowing Gal4 to trigger FLP expression and facilitating FLP/*FRT* recombination within the *esg-Gal4* lineage, which includes ISCs. We were thus able to induce *Ras* oncogenic mutations in all cell types in the adult midgut (Figure 3B). Following induction, we maintained adult flies at 18°C and analyzed their intestinal barrier function using the Smurf assay (Figure 3C)^36^. This simple blue dye feeding assay detects intestinal barrier dysfunction by causing the entire body to turn blue. As expected, we did not observe blue coloration in *esg-Gal4/+; tub-Gal80^ts^, UAS-FLP/Ras85D^GFP>>wt^* control animals (Figure 3D). In contrast, defects in barrier function were observed in our *Ras* allele switch oncogenic lines. Surprisingly, unlike the effects observed in the wing disc, *esg-Gal4/+; tub-Gal80^ts^, UAS-FLP/Ras85D^GFP>>G12V^* animals exhibited significantly more pronounced intestinal barrier defects than were observed in *esg-Gal4/+; tub-Gal80^ts^, UAS-FLP/Ras85D^GFP>>Q61H^* flies (Figure 3D). These results suggest that the *Ras G12V* mutation has a more detrimental impact on the adult midgut than the *Ras Q61H* mutation, highlighting the tissue-dependent effects of oncogenic mutations.

**Figure 3.**
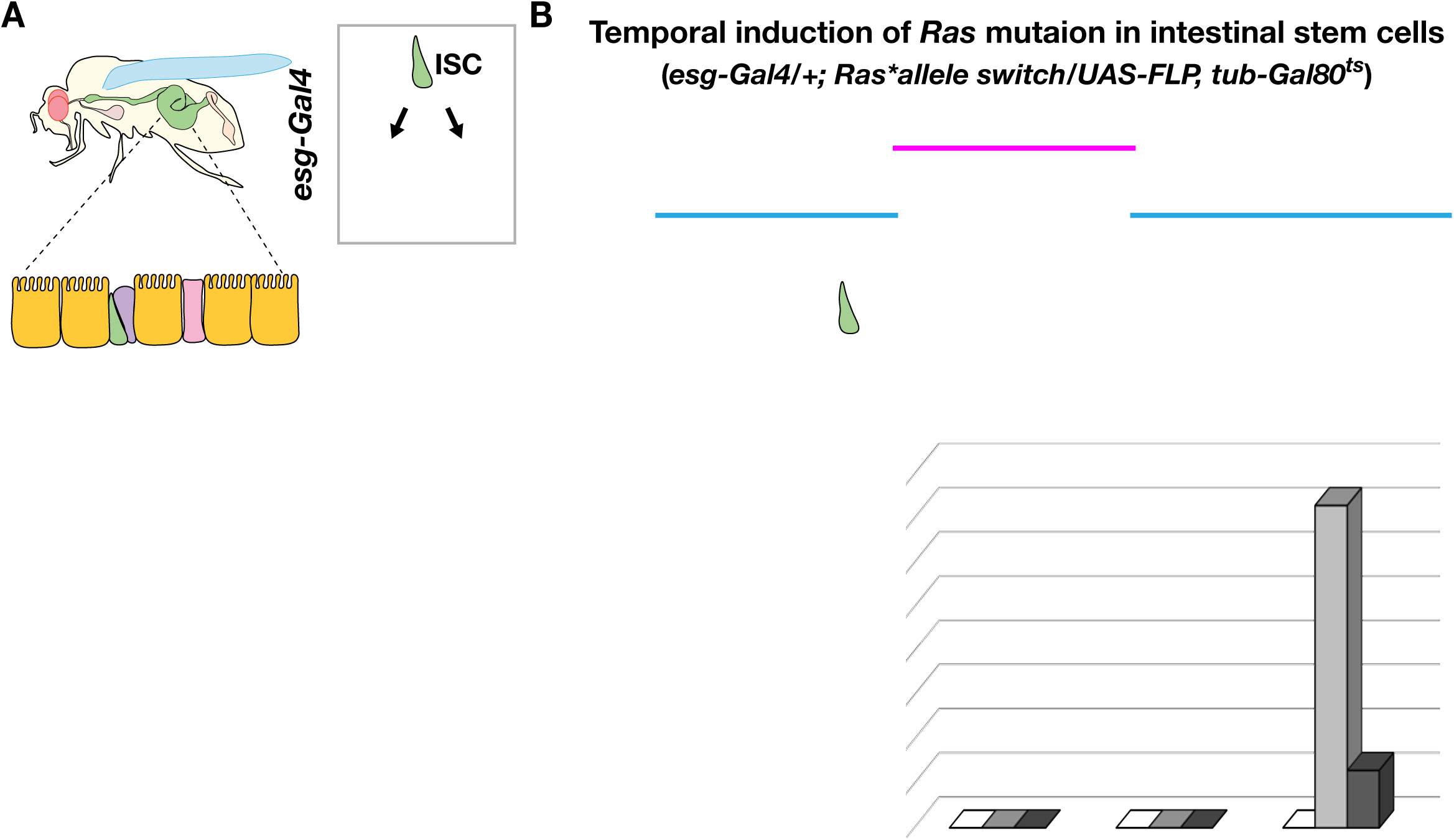
*Ras G12V* causes intestinal barrier dysfunction. (**A**) *Drosophila* intestinal stem cells (ISCs) differentiate into enteroblasts (EBs) and enteroendocrine progenitors (EEPs), which eventually develop into enterocytes (ECs) and enteroendocrine cells (EEs). *esg-Gal4* is active in ISCs, EBs and EEPs. (**B**) An experimental design for temporally inducing *Ras* oncogenic mutations in ISCs. Animals were reared at 18°C during development, and 2-day old adult flies were moved to a 29°C incubator to induce *Ras85D* oncogenic mutations. After 2-days induction, flies were kept at 18°C and were collected at indicated time points to check their intestinal function. (**C**) A schematic illustration of the Smurf assay. Flies are fed with food containing blue dye overnight. Flies turn blue if they have intestinal epithelial barrier problems. (**D**) Quantification of the Smurf assay. Intestinal epithelial barrier defect was not observed in *esg-Gal4/+; Ras85D^GFP>>wt^/UAS-FLP, tub-Gal80^ts^* control flies (10-day: *n*=790, 20-day: *n*=720, 30-day: *n*=624). Flies expressing *Ras G12V* exhibit more Smurf phenotypes than those with the *Ras Q61H* mutation. *esg-Gal4/+; Ras85D^GFP>>G12V^/UAS-FLP, tub-Gal80^ts^* (10-day: *n*=843, 20-day: *n*=769, 30-day: *n*=740). *esg-Gal4/+; Ras85D^GFP>>Q61H^/UAS-FLP, tub-Gal80^ts^* (10-day: *n*=516, 20-day: *n*=364, 30-day: *n*=307).

### Induction *of Ras Q61H* causes progressive clonal regression

To investigate the cellular mechanisms underlying the tissue-dependent effects of *Ras* oncogenic alleles, we first determined whether the *Ras G12V* and *Q61H* mutations influenced cell proliferation in the adult midgut (Figure 4). Ten days after the *Ras85D* allele switch, we analyzed adult midguts stained with anti-p-H3 antibodies from control (*esg-Gal4/+; tub-Gal80^ts^, UAS-FLP/Ras85D^GFP>>wt^*) and experimental animals (*esg-Gal4/+; tub-Gal80^ts^, UAS-FLP/Ras85D^GFP>>G12V^* or *Ras85D^GFP>>Q61H^*). Interestingly, despite the previously observed intestinal barrier dysfunction, cell clones expressing the *Ras G12V* mutation did not exhibit a statistically significant difference in the number of mitotic cells compared to controls (Figure 4A-D and G). In contrast, we found that the *Ras Q61H* mutation significantly increased cell proliferation, leading to intestinal hyperplasia 10 days after the induction (Figure 4E-G).

**Figure 4.**
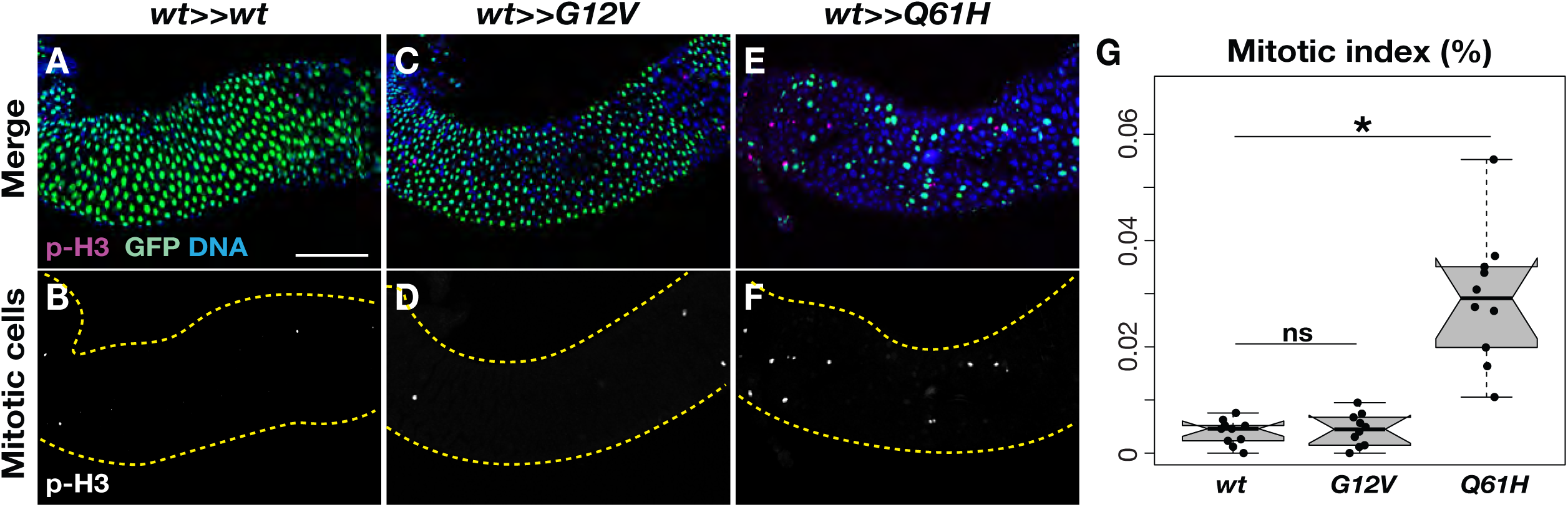
*Ras Q61H* promotes cell division in the adult midgut. **(A-F)** Mitotic cells in the adult midguts were detected 10 days after the *Ras85D* allele switch by anti-p-H3 staining (*magenta*). The panels **B**, **D**, and **F** only display p-H3 positive cells. *Yellow* dashed lines indicating the outlines of the midguts. Scale bar: 100 μm. GFP (*green*) indicates cells expressing *wild-type Ras85D*, while the mutant cells lost GFP signals. **(G)** Mitotic index of the control *esg-Gal4/+; tub-Gal80^ts^, UAS-FLP/Ras85D^GFP>>wt^*, *esg-Gal4/+; tub-Gal80^ts^, UAS-FLP/Ras85D^GFP>>G12V^*, *esg-Gal4/+; tub-Gal80^ts^, UAS-FLP/Ras85D^GFP>>Q61H^* midguts (*n*=10 for each genotype). Two-sided Student’s *t-test*: \**P* < 0.05, not significant (ns; *P* > 0.05).

To investigate the temporal dynamics of *Ras* mutant clone persistence in the midgut, we next collected samples at 0, 5, 10, 20, and 30 days after induction using the *esg^ts^FLP* system. We then quantified the number of mutant cells per clone over time (Figure 5). Although the *Ras G12V* mutant clones did not show an observable increase in cell proliferation compared to controls, mutant cells in *esg-Gal4/+; tub-Gal80^ts^, UAS-FLP/Ras85D^GFP>>G12V^* midguts nevertheless steadily increased in number, progressively occupying almost the entire midgut by day 30 (Figure 5A-L). In contrast, *Ras Q61H* mutant cells increased in number rapidly and occupied nearly the entire midgut by day 10, consistent with a higher rate of proliferation (Figure 5M-O, and R). Strikingly, after this rapid clonal expansion the number of *Ras Q61H* mutant cells then gradually regressed over time, allowing the GFP-positive control cells to repopulate the *esg-Gal4/+; tub-Gal80^ts^, UAS-FLP/Ras85D^GFP>>Q61H^* midguts (Figure 5O-R). Since MAPK signaling plays a critical role in Ras-dependent cell proliferation, we employed a *Raf* null mutant, *Raf*^7^ to test for effects of cell proliferation on the behavior of *Ras Q61H* mutant clones. In a *Raf* heterozygous background, *Ras Q61H* mutant cells in the midguts propagated more slowly and persisted longer compared to those in a *wild-type* background (Figure 5M-X; *Raf^7^/+; esg-Gal4/+; tub-Gal80^ts^, UAS-FLP/Ras85D^GFP>>Q61H^*). These findings suggest that the increased proliferative capacity of *Ras Q61H* clones contributes to their eventual regression, and that MAPK signaling levels play a critical role in regulating the dynamic behaviors of *Ras* mutant cells.

**Figure 5.**
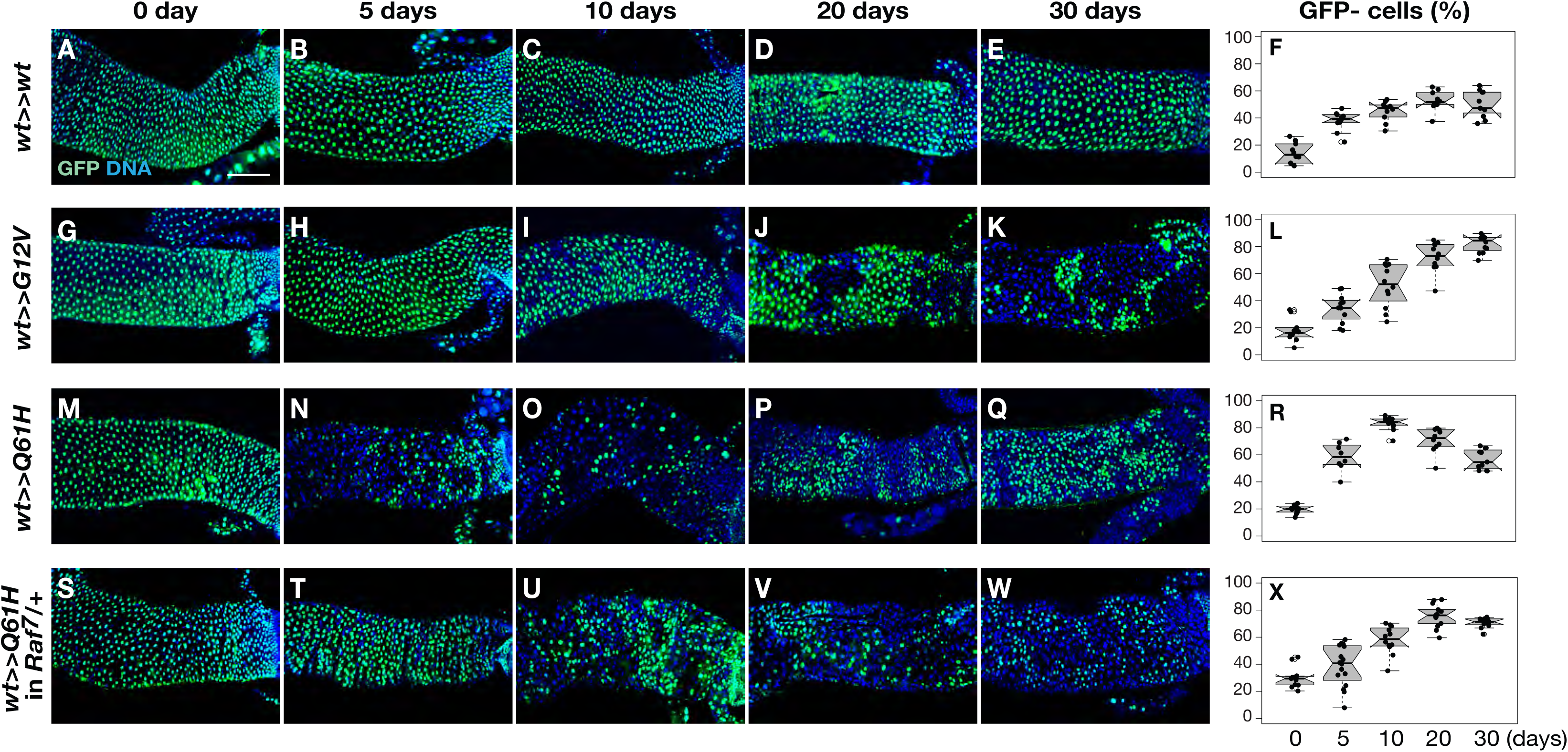
*Ras Q61H* leads to the clonal regression of mutant cells in the adult midgut. **(A-X)** After inducing *Ras85D* allele switch by temperature shift, midguts were collected from control *esg-Gal4/+; tub-Gal80^ts^, UAS-FLP/Ras85D^GFP^*^>*>wt*^ (**A-F**, 0-day: *n*=16, 5-day: *n*=10, 10-day: *n*=10, 20-day: *n*=10, 30-day: *n*=11), *esg-Gal4/+; tub-Gal80^ts^, UAS-FLP/Ras85D^GFP>>G12V^* (**G-L**, 0-day: *n*=10, 5-day: *n*=11, 10-day: *n*=12, 20-day: *n*=12, 30-day: *n*=12), *esg-Gal4/+; tub-Gal80^ts^, UAS-FLP/Ras85D^GFP>>Q61H^*(**M-R**, 0-day: *n*=10, 5-day: *n*=8, 10-day: *n*=12, 20-day: *n*=10, 30-day: *n*=11), and *Raf^7^/+; esg-Gal4/+; tub-Gal80^ts^, UAS-FLP/Ras85D^GFP>>Q61H^* midguts (**S-X**, 0-day: *n*=11, 5-day: *n*=16, 10-day: *n*=12, 20-day: *n*=14, 30-day: *n*=12). Loss of GFP (g*reen*) signals represents cells expressing different *Ras85D* alleles due to the allele switch. Scale bar: 100 μm. The number of GFP-negative cells was quantified in **F**, **L**, **R**, and **X**.

## Discussion

Oncogenic mutations have dominant effects, and thus, functional studies of these mutations have primarily relied on overexpression analysis^22, 37^. While overexpression systems like Gal4/UAS are convenient and powerful genetic tools, there are important conceptual limits to their utility. Exogenous gene expression approaches tend to push transgene expression to non-physiological levels, overstimulating the biological system and potentially obscuring the more subtle characteristics of specific alleles. To rigorously examine the effect of *Ras* mutations, here we developed several inducible *Ras85D* oncogenic lines by modifying the endogenous *Ras* locus using CRISPR/Cas9-mediated genome engineering (Figure 1D-H; Figure S3). Contrasting with ectopic oncogene expression driven by exogenous regulatory elements not subject to feedback control, our inducible *Ras85D* mutant lines permit the induction of single copy oncogenic alleles within living tissues under the endogenous gene regulatory circuitry (Figure 1D-H; Figure S3). Thus, our inducible lines are not only ideal to accurately assess oncogenic activity *in vivo* but also suggest a new platform for exploring upstream and downstream regulators of the *Ras* gene. Further, the founder line for the *Ras85D* allele switch enables the creation of new oncogenic inducible lines via simple PhiC31 integrase-mediated transformation, facilitating analysis for additional alleles (Figure S3A).

Errors in DNA replication during cell division are a major cause of cancer^38^. Most oncogenic mutations contribute to cancer development by increasing the chances of acquiring additional mutations, as they accelerate cell proliferation and promote the expansion of clonal populations^15–19^. Interestingly, our findings revealed that endogenous *Ras* oncogenic mutations did not impact the rate of cell proliferation in the wing disc (Figure 2A-D). Instead, these mutations, except for *Ras G12V*, induced the formation of cysts and disrupted epithelial architecture (Figure 2E-H; Figure S5). Previous studies have shown that apical mitotic rounding is critical for creating sufficient physical space for spindle formation and chromosome segregation, thus ensuring proper mitosis^39, 40^. Chromosome missegregation often leads to cell death but can also result in viable aneuploid cells which in turn contribute to tumorigenesis^41, 42^. In the present study, the abnormal epithelial architecture of cysts may impose physical constraints on mitotic cells, leading to a microenvironment that enhances the likelihood of chromosome segregation defects. Consistent with this, we frequently observed cell death associated with extruding *Ras* mutant cysts (Figure 3L).

In contrast to the defects observed in the developing wing, *Ras* oncogenic mutations exhibited minimal impact on the structure of the adult midgut. *Ras* mutant cells differentiated into EEs and ECs, and most midguts carrying mutant cells maintained epithelial integrity (Figure 3D; Figure S6). However, consistent with the conventional view, *Ras* oncogenic mutations conferred a growth advantage that led to hyperplasia as previously described (Figure 4; Figure 5)^37^. These results suggest that the primary effect of *Ras* oncogenic mutations in the adult midgut is to increase the likelihood of acquiring additional mutations due to DNA replication errors. In sum, our findings indicate two distinct molecular actions of *Ras* oncogenic mutations that could theoretically contribute to cancer development, highlighting a potentially critical role for aberrant tissue biomechanics in oncogenesis.

Cellular and phenotypic plasticity are crucial factors in the initiation and progression of cancer^43–45^. The first step in cancer development involves the hyper-proliferation of tumor cells. Interestingly, our study revealed unexpected effects of oncogenic allele-dependent cellular behaviors on the expansion of tumor cells in the adult midgut. While *Ras G12V* mutant cells consistently increased in population over time, *Ras Q61H* mutant cells underwent regression after a rapid expansion phase (Figure 5A-R). Our findings suggest that allele-dependent cellular behaviors may contribute to the cancer type-specific accumulation of *Ras* oncogenic mutations (Figure 6). It has been proposed that cell fitness-based selection plays a key role in tumor cell expansion^46, 47^. For instance, previous studies using a mammalian epithelial *in vitro* model and the *Drosophila* wing disc demonstrated that *Ras* mutant cells are apically extruded when existing as mosaics^48, 49^. In addition, in mouse skin epithelia, injury stimulates the elimination of *Ras* mutant cells by *wild-type* cells, thereby preventing tumor cell expansion^50^. These results suggest that tumor cell regression could be a common protective mechanism, playing a critical role in the early stages of cancer development. Moreover, in the presence of a heterozygous *Raf* null mutation, we found that *Ras Q61H* mutant cells persisted longer than in the *wild-type* genetic background (Figure 5M-X). This result implies that the regression process is regulated in a MAPK activity-dependent manner. Given that many oncogenic mutations occur within the RTK/Ras/MAPK pathway^51, 52^, it would be of great interest to investigate whether mutations in the components of this signaling pathway cause similar tumor cell dynamics. Lastly, building on our findings, we speculate that a partial reduction of MAPK activity through any means, such as drug treatments, germline genotypes, or environmental factors, could prolong the persistence of tumor cells and thus increase the likelihood of acquiring additional deleterious mutations in some tissue contexts.

**Figure 6.**
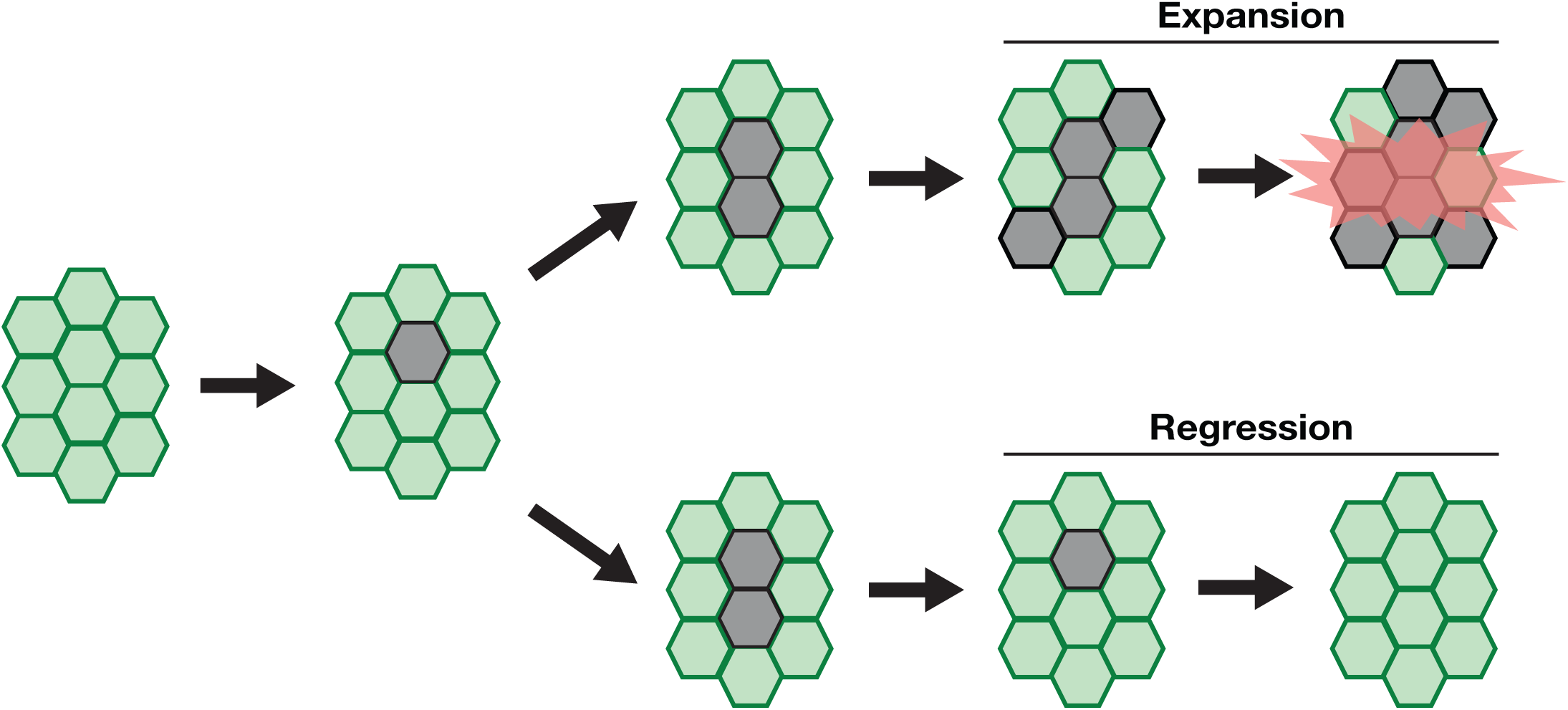
Allele- and tissue- dependent *Ras* oncogenic activity. Different tumor cell behaviors contribute to the tissue-specific accumulation of *Ras* oncogenic mutations.

## Materials and methods

### *Drosophila* maintenance and stocks

Flies were reared using standard cornmeal, molasses and yeast media at 25°C, except where noted. The fly stocks used in this study were *y^1^, vas-Cas9 ZH-2A* (BDSC#66554), *y^1^ M{RFP[3xP3.PB] GFP^E.3xP3^=vas-int.Dm}ZH-2A w\** (BDSC#40161), *w; nubbin-Gal4/CyO; UAS-FLP*^53^, *w hs-flp* (BDSC#8862), *esg-Gal4/CyO sChFP; UAS-FLP3, tub-Gal80ts/TM6C* (this study), *puc-lacZ*(*P{ry[+t7.2]=A92}puc[E69]/TM3, Sb[1]*, BDSC#98329), and *Raf^7^/FM7a* (BDSC#7338). We also established six *Ras85D* allele switch transgenic lines, as we described in detail below.

### Generation of inducible *Ras85D* allele transgenic lines

All primers used to generate inducible *Ras85D* allele transgenic lines were listed in the Supplementary Table. To generate two *Ras85D* sgRNA DNA constructs, primers 1 - 4 were annealed and cloned into the BbsI site of a sgRNA expression vector *pBFv-U6.2*^54^. Next, we prepared a *Ras85D* donor DNA construct with a ubiquitin promoter mCherry selection maker in pHSG298 backbone^53, 55^. Five DNA fragments were amplified using primers 5 to 14. After the PCR reaction, five PCR products were purified using Zymoclean Gel DNA Recovery Kit (D4001, Zymo Research) and combined using HiFi DNA Assembly Master Mix (E2621S, NEB). To establish a *Ras85D* allele switch founder line, we injected two sgRNA DNAs and the donor plasmid (250 ng/ml for each) into the posterior side of *vas-Cas9 ZH-2A* embryos. Transgenic flies were screened using the ubiquitous expression of mCherry. mCherry-positive candidate lines were further molecularly confirmed by PCR and DNA sequencing.

To prepare *Ras85D* cassette DNAs, we first generated a *w+attB ubi-GFP* vector. A ubiquitin promoter-nuclear eGFP fragment with 5’ SalI and 3’ MluI was obtained from *pHSG298 dpp donor ubi-GFP* using primer primers 15 and 16^56^. Then, we cloned the *ubi-GFP* fragment into XhoI and MluI sites of *w+attB* (#30326, addgene). Second, a *Ras85D* DNA fragment was amplified using primers 17 and 18 with 5’ MluI and 3’ AscI, and the resulting PCR product was cloned into *pCRII blunt TOPO* (K280002, Thermo Fisher Scientific). Each *Ras85D* oncogenic mutation was induced with the Q5 Site-Directed Mutagenesis Kit (E0554S, NEB), using primers 19 to 27. After DNA sequencing, *pCRII blunt TOPO Ras85D* DNAs with desired oncogenic mutations were digested by MluI and AscI and cloned into *w+attB ubi-GFP*.

To establish *Ras85D* allele switch transgenic lines, *w+attB ubi-GFP Ras85D* DNA constructs carrying different oncogenic mutations were injected into the posterior regions of embryos obtained from a cross of *y^1^ M{RFP[3xP3.PB] GFP^E.3xP3^=vas-int.Dm}ZH-2A w\** (BDSC#40161) and the *w*; *Ras85D* allele switch founder. Transformants were screened by the presence of red eye color, and the selection markers were removed via Cre/*loxP*-mediated recombination (Figure S3A).

### Drosophila crosses

Induction of *Ras* oncogenic alleles in the wing pouch region (Figure 1E-H; Figure 2A-D; Figure S3B-S)

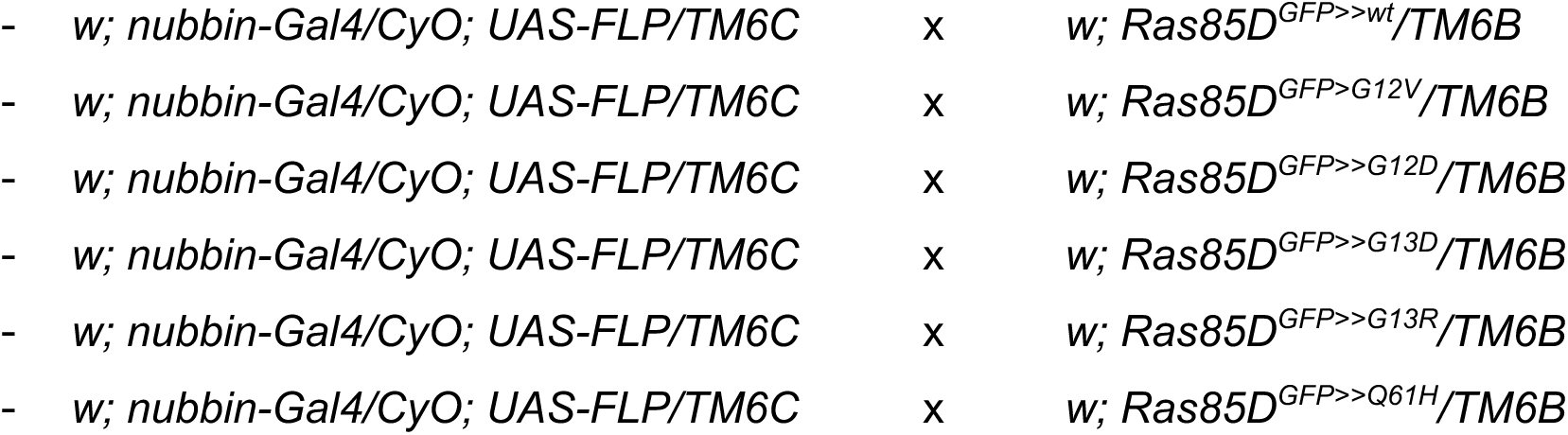

Random induction of *Ras* oncogenic alleles in the wing disc (Figure 2F-L; Figure S4; Figure S5)

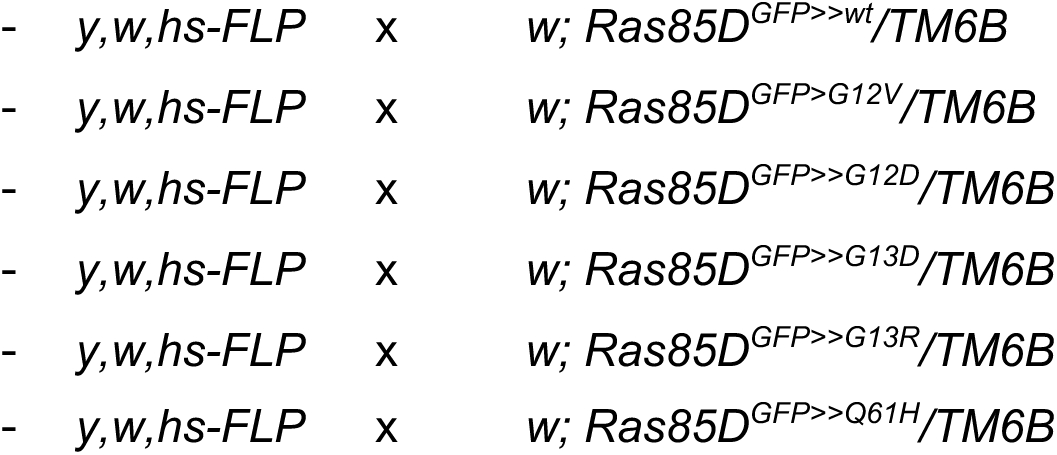

Induction of *Ras* oncogenic alleles in the adult midgut (Figure 3D; Figure 4; Figure 5; Figure S6)

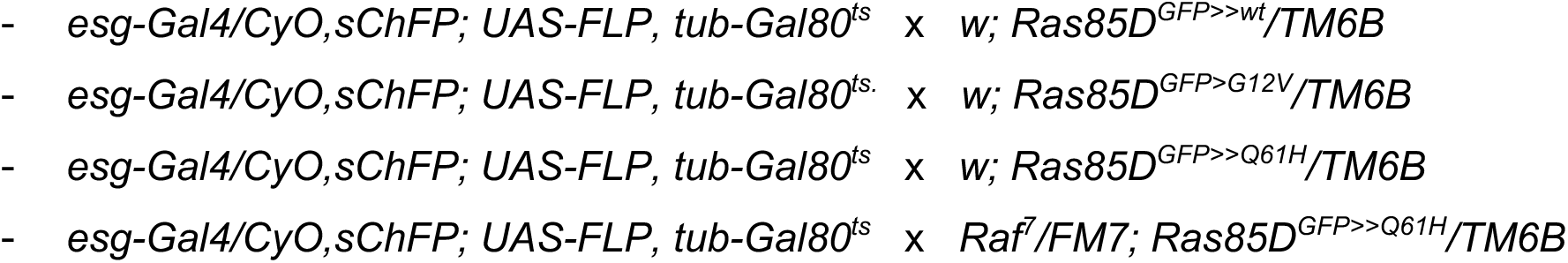

### Immunohistochemistry and imaging

Female third-instar wing discs and adult midguts were dissected in PBS (137 mM NaCl, 27 mM KCl, 100 mM Na_2_HPO_4_, and 18 mM KH_2_PO_4_, pH 7.4), fixed with 4% PFA/PBS at room temperature for 20 minutes. After three PBST (PBS + 0.1% Triton X-100) washes at room temperature for 20 minutes, samples were blocked using PBST + 10% normal goat serum at room temperature for 1 hour followed by primary antibody incubation at 4°C overnight. Primary antibodies used in this study were rabbit anti-dp-ERK (1:200; #4370, Cell Signaling Technology), mouse anti-*Drosophila* Dlg (1:500; 4F3, DSHB), mouse anti-p-H3 (1:1,000; #05-806, Millipore Sigma), mouse anti-β-galactosidase (1:200; Z3781, Promega) and rabbit anti-cleaved *Drosophila* Dcp1 (1:500; Asp216, Cell Signaling Technology). After overnight incubation, tissue samples were washed with PBST three times at room temperature for 20 minutes and were incubated with Alexa Fluor conjugated secondary antibodies (1:500, Thermo Fisher Scientific) and Hoechst 33342 (1:1,000, H3570, Thermo Fisher Scientific) at room temperature for 3 hours. Actin was visualized by using SiR-Actin (1:1,000, CY-SC001, Cytoskeleton). All primary and secondary antibodies and staining reagents were diluted in PBST. After three PBST washes at room temperature for 20 minutes, samples were mounted using VECTASHIELD Antifade Mounting Medium (H-1000-10, Vector Laboratories). All images were obtained using a Leica TCS SP5, a Leica TCS SP8, or a Zeiss LSM780 confocal microscopy.

### Imaging analyses

The mitotic index was calculated by dividing the number of mitotic cells (identified by p-H3 positivity) by the total number of cells (visualized by anti-Dlg or Hoechst 33342 staining) using a 30.03 μm square in the anterior–ventral wing pouch region and the posterior midgut. The ‘find maxima’ function of FIJI was used to automatically count total cell numbers.

## Acknowledgements

We thank the Bloomington Stock Center for fly stocks and the Developmental Studies of Hybridoma Bank for anti-Dlg antibody. We are grateful to members of the Gibson lab for discussion and advice. We also thank Abby Dreyer at the Stowers Institute for Medical Research and Joie Harney and Alicia Thome at Indiana State University for their administrative support. This work was supported by funding from the Stowers Institute for Medical Research, NIH R01 GM111733 (M.C.G.), Indiana State University, and Indiana Academy of Science Senior Research Grants IASSG-F23-01 (T.A.).

## Author Contribution

T.A. and M.C.G. conceived the project, designed the experiments and wrote the manuscript. T.A. performed the experiments and analyzed the data.

---

**Figure S1.**
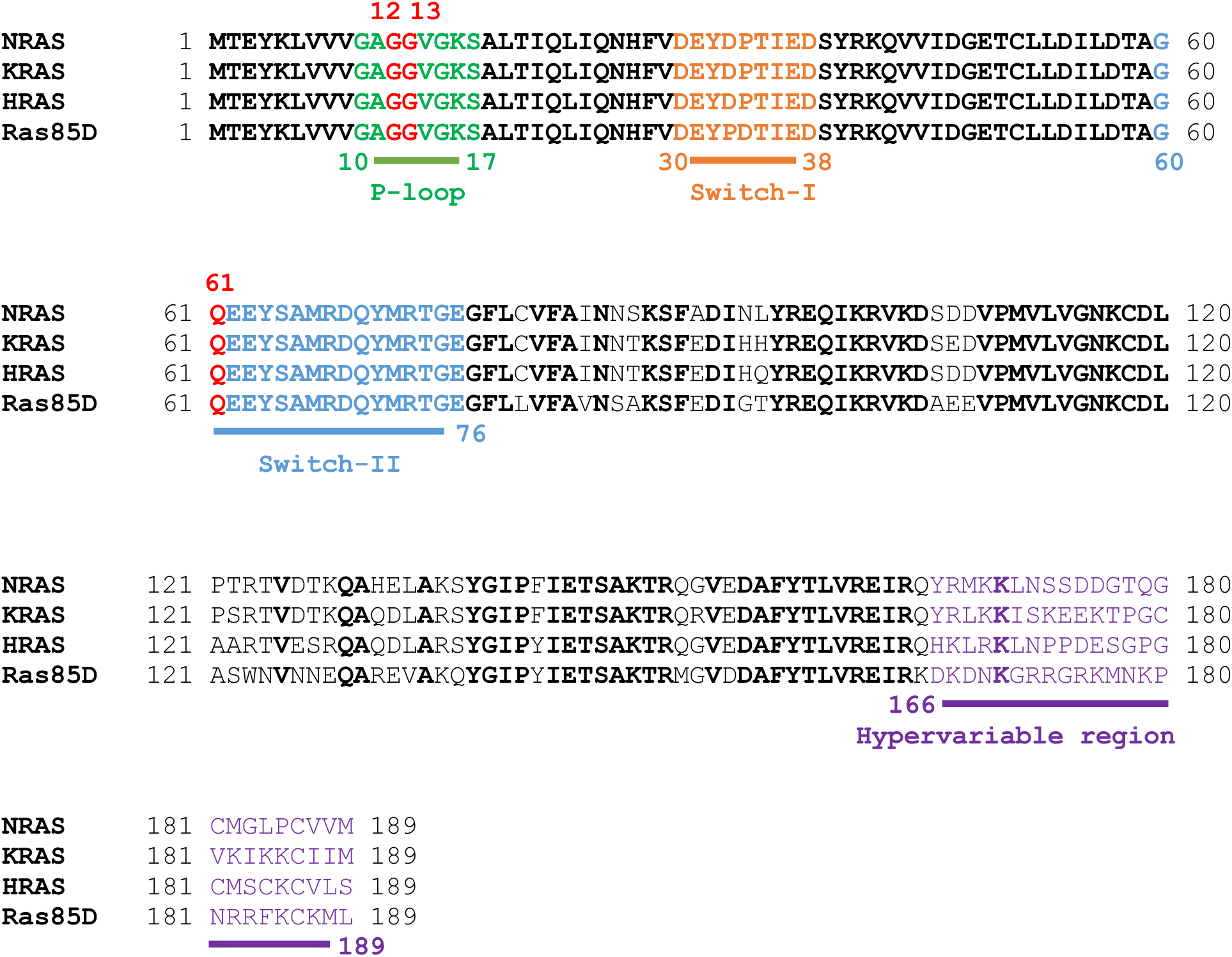
Comparison between human K-Ras, N-Ras, H-Ras and *Drosophila* Ras85D proteins. Oncogenic mutation hot spots (G12, G13 and Q61) and each protein domain are highlighted. The bold characters indicate the conserved amino acid residues between three Ras proteins.

**Figure S2.**
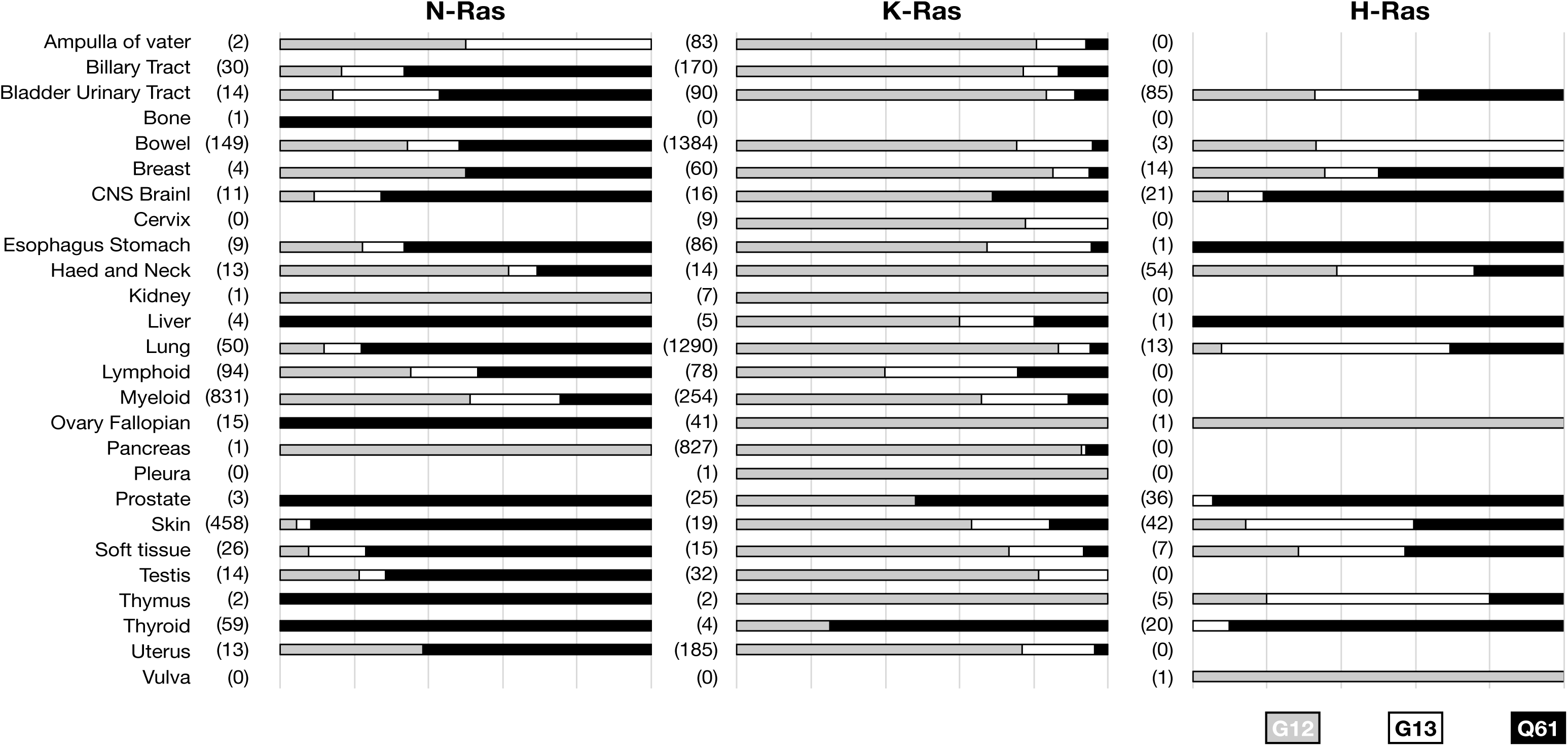
Tissue-specific accumulation of *K-Ras*, *N-Ras*, and *H-Ras* oncogenic mutations. The numbers in parentheses indicate sample sizes.

**Figure S3.**
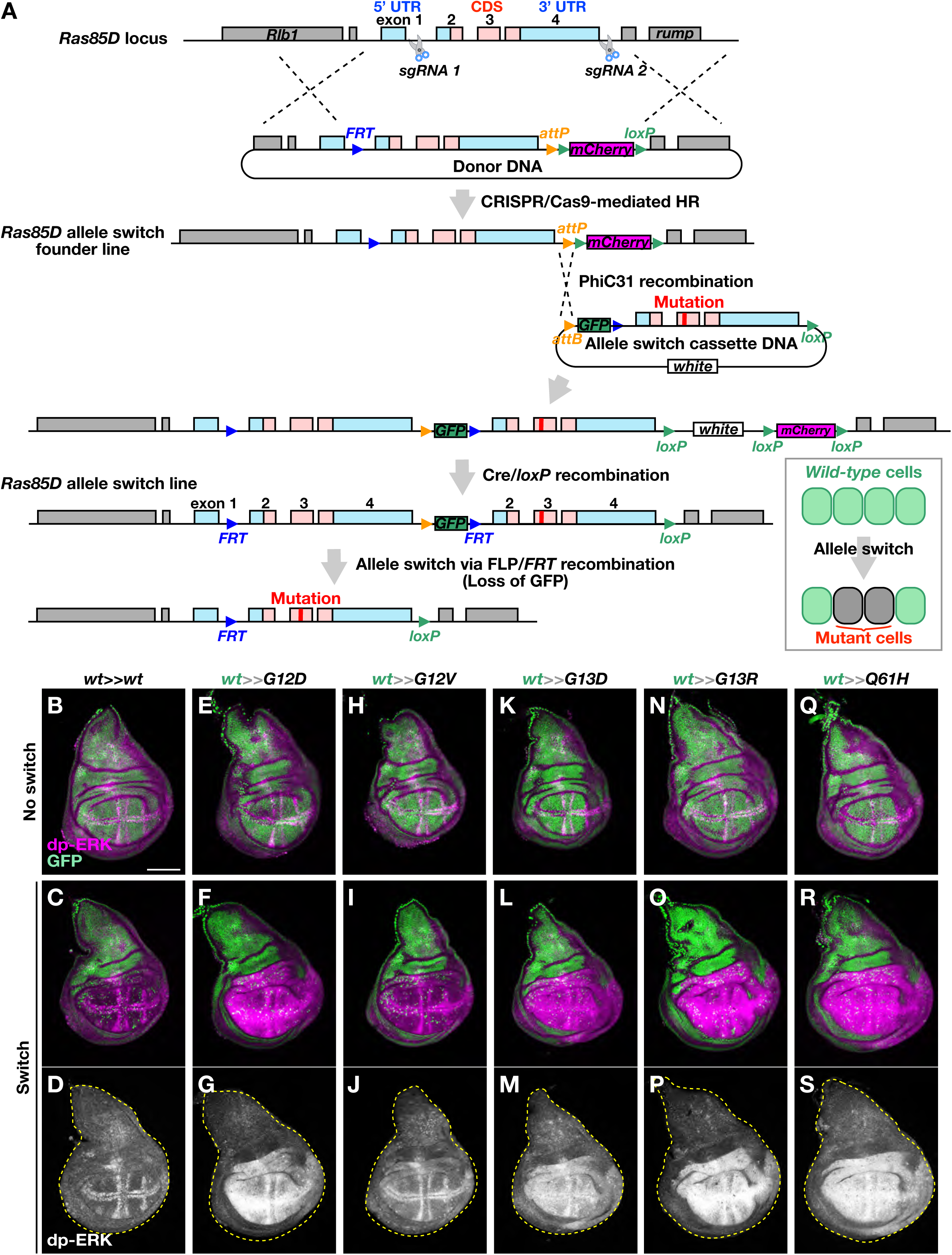
Generation of *Ras85D* inducible alleles. **(A)** The schematic illustration depicts the process of establishing inducible *Ras85D* alleles. **(B-S)** Different *Ras85D* alleles were induced in the wing pouch area using *nubbin-Gal4* and *UAS-FLP*. The activation of the MAPK pathway was visualized through anti-dp-ERK staining (*magenta*) in the wing discs, both before **(B**, **E**, **H**, **K**, **N**, **Q)** and after **(C**, **D**, **F**, **G**, **I**, **J**, **L**, **M**, **O**, **P**, **R**, **S)** the allele switch induction. Loss of GFP (*green*) signals indicates the population of cells switched from *wild-type* to desired *Ras85D* alleles. Scale bar: 100 μm.

**Figure S4.**
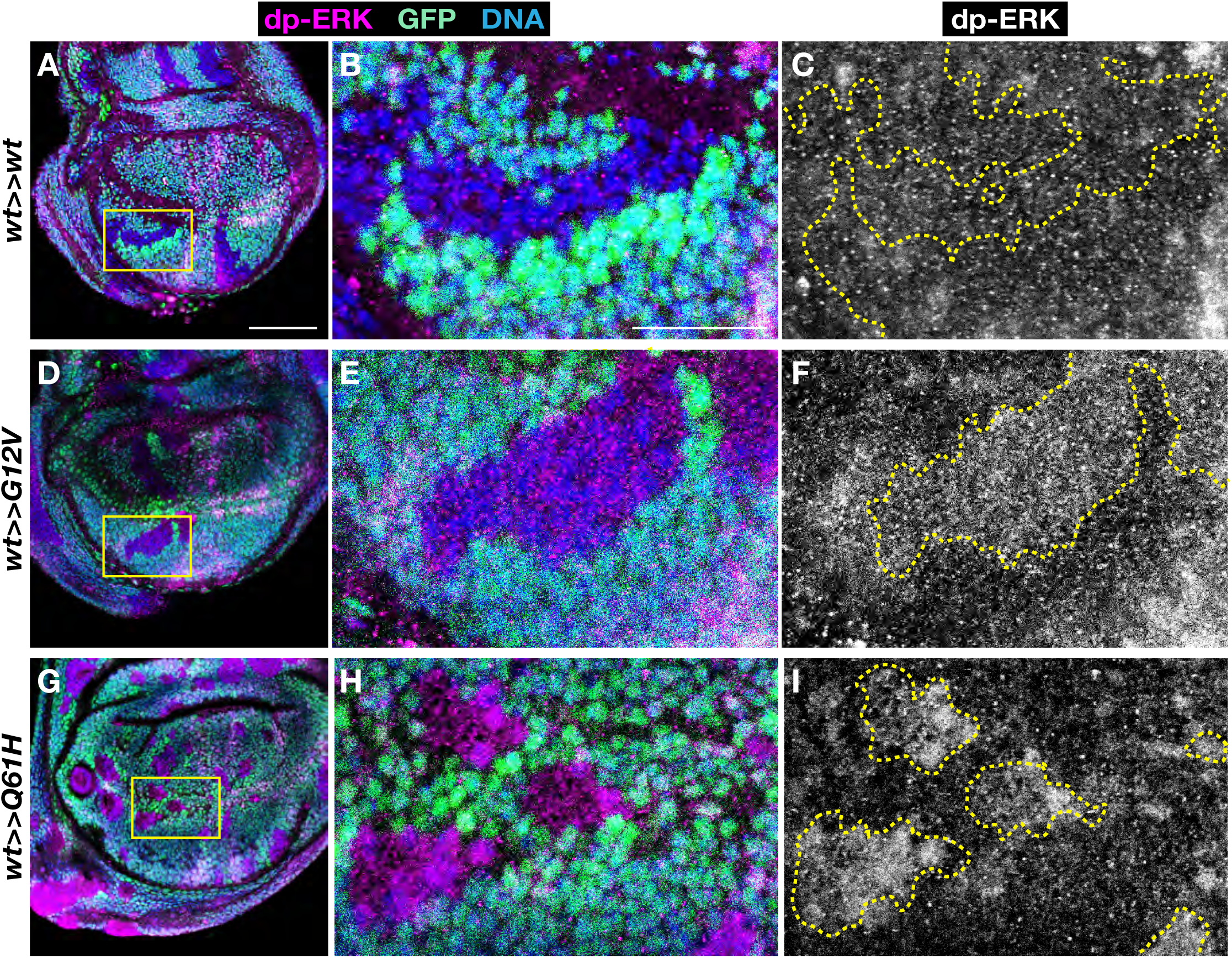
*Ras85D* oncogenic mutation triggers a cell-autonomous activation of MAPK signaling. **(A-I)** Small clones of cells expressing *wild-type Ras85D* (**A**-**C**), *G12V* (**D**-**F**), and *Q61H* (**G**-**I**) were randomly induced in the wing disc using *hs-FLP*. dp-ERK staining demonstrated a cell-autonomous MAPK activation in the clones of cells expressing *Ras85D* oncogenic mutations marked by loss of GFP (*green*). 50 μm in **A**; 20 μm in **B.**

**Figure S5.**
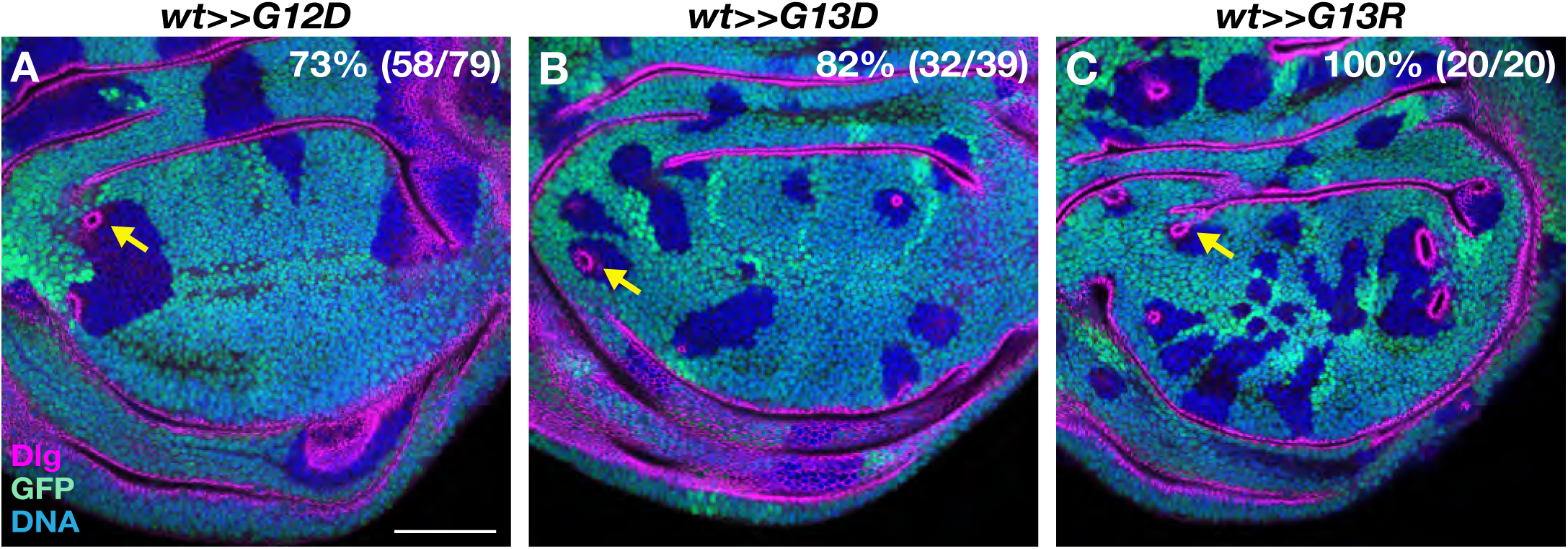
*Ras85D* oncogenic *G12D*, *G13D*, and *G13R* mutations lead to the cyst formation the wing disc. **(A-C)** *G12D* (**A**), *G13D* (**B**), and *G13R* (**C**) oncogenic mutations induced cysts in the wing discs. The locations of mutant clones are indicated by the loss of GFP signals (*green*). Scale bar: 50 μm.

**Figure S6.**
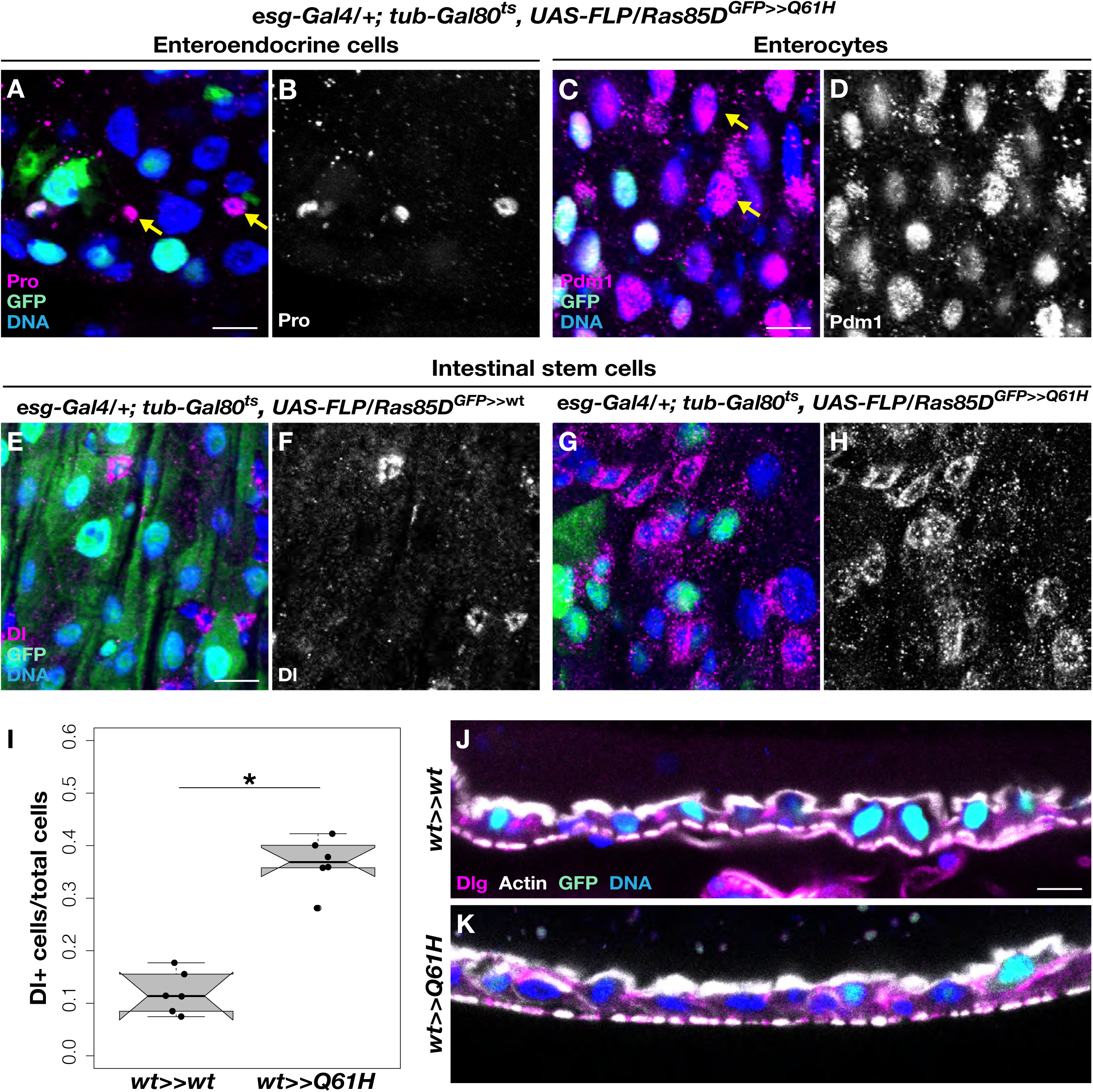
*Ras Q61H* mutant cells differentiate into enteroendocrine cells and enterocytes, maintaining the integrity of the epithelial structure. (**A**-**D**) *Ras Q61H* mutant cells in the *esg-Gal4/+; tub-Gal80^ts^, UAS-FLP/Ras85D^GFP>>Q61H^* midguts generate Prospero positive enteroendocrine cells (Pros, *magenta* in **A**) and enterocytes expressing Pdm1 (*magenta* in **C**). (**E-I**) Inducing *Ras* Q61H mutations using *esg^ts^FLP* increased the intestinal stem cells detected by anti-Delta antibody compared to the control midguts (Dl, *magenta* in **E** and **G**). (I) Quantification of intestinal stem cell numbers. Two-sided Student’s *t-test*: \**P* < 0.05. (**J**-**K**) Intestinal epithelial structures of the control *esg-Gal4/+; tub-Gal80ts, UAS-FLP/Ras85D^GFP>>wt^* and *esg-Gal4/+; tub-Gal80ts, UAS-FLP/Ras85D^GFP>>Q61H^* midguts visualized by anti-Dlg and Sir-Actin staining. Scale bars: 10 μm.

**Supplementary Table.**
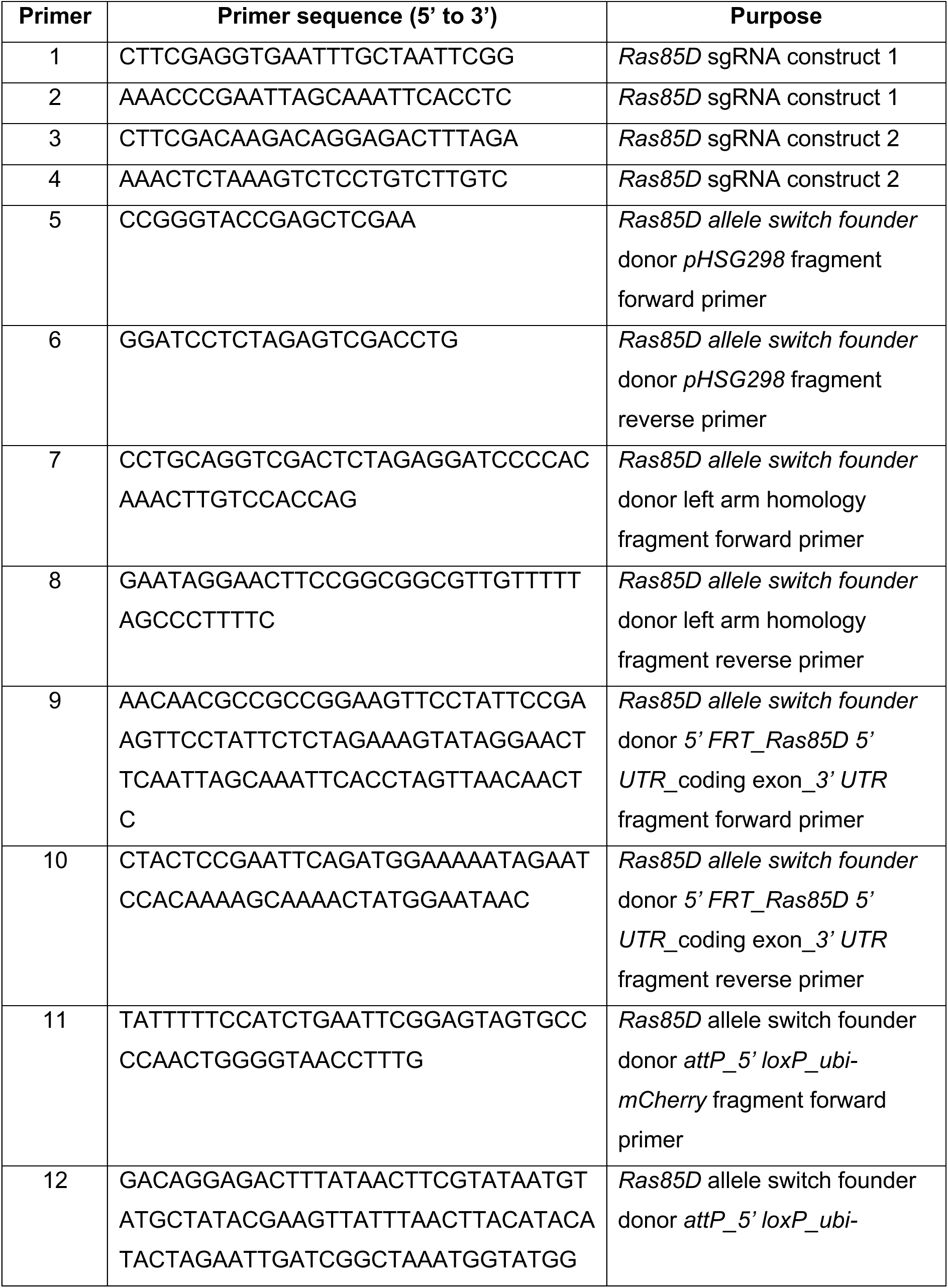
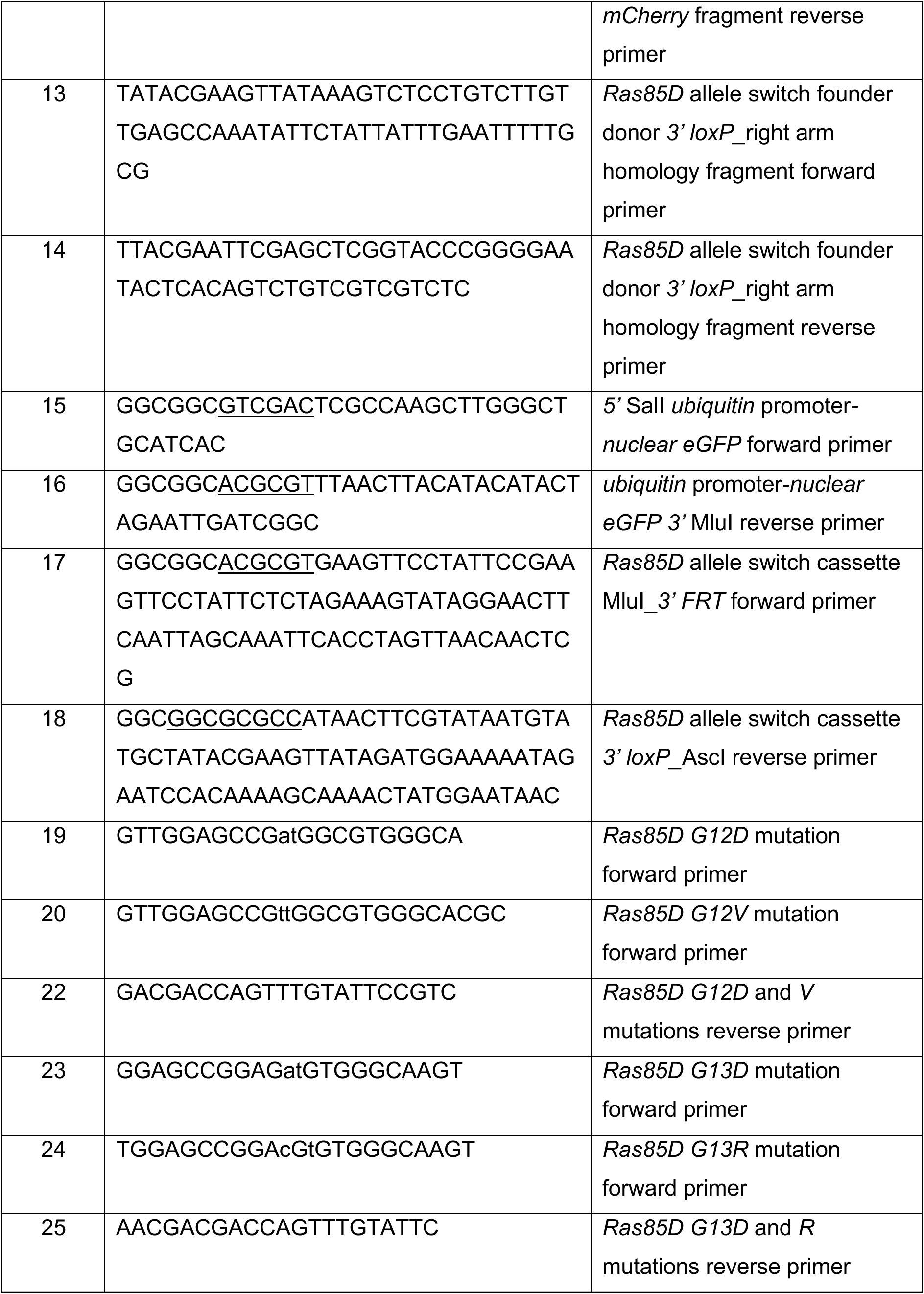
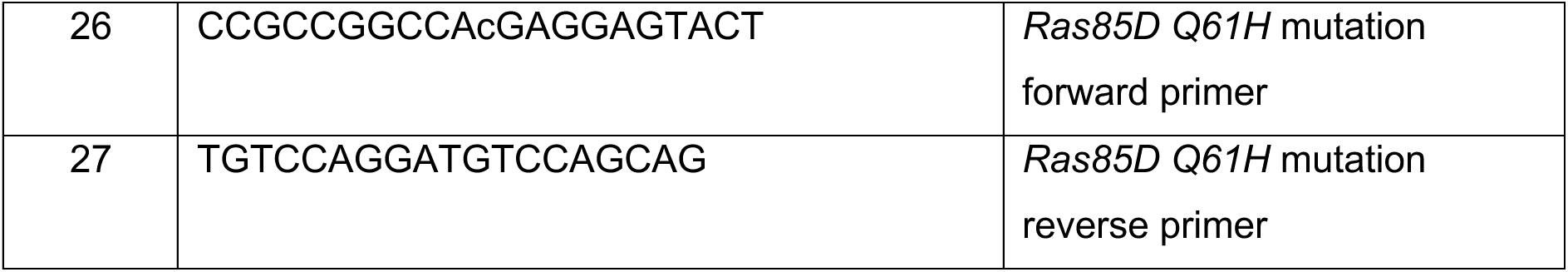
Primers used in this study. Restriction enzyme sites are underlined in primer sequences. Lowercase characters indicate point mutations.

